# Carbon nanotube electrodes for retinal implants: a study of structural and functional integration over time

**DOI:** 10.1101/050633

**Authors:** Cyril G. Eleftheriou, Jonas B. Zimmermann, Henrik D. Kjeldsen, Moshe David-Pur, Yael Hanein, Evelyne Sernagor

## Abstract

The choice of electrode material is of paramount importance in neural prosthetic devices. Electrodes must be biocompatible yet able to sustain repetitive current injections in a highly corrosive environment. We explored the suitability of carbon nanotube (CNT) electrodes to stimulate retinal ganglion cells (RGCs) in a mouse model of outer retinal degeneration. We investigated morphological changes at the bio-hybrid interface and changes in RGC responses to electrical stimulation following prolonged *in vitro* coupling to CNT electrodes. We observed gradual remodelling of the inner retina to incorporate CNT assemblies. Electrophysiological recordings demonstrate a progressive increase in coupling between RGCs and the CNT electrodes over three days, characterized by a gradual decrease in stimulation thresholds and increase in cellular recruitment. These results provide novel evidence for time-dependent formation of viable bio-hybrids between CNTs and the retina, demonstrating that CNTs are a promising material for inclusion in retinal prosthetic devices.

## Introduction

Retinal prostheses aim to restore a degree of vision in patients with photoreceptor degeneration. The principle is either to take advantage of the surviving retinal circuitry or to target retinal ganglion cells (RGCs) directly, as these cells are the only output from the eye to the brain, encoding visual scenes into spike trains which are then transmitted to central visual targets via the optic nerve^1, 2^. Epi-retinal prostheses consist of micro-electrode arrays (MEAs) apposed to the vitreal side of the retina, providing direct electrical stimulation to the RGC layer^3^. Individual electrodes have to deliver electrical pulses strong enough to elicit action potentials in surrounding RGCs without damaging the electrode material or the target tissue^4^.

As the main interface between the prosthetic device and the tissue, electrodes are critically important components of any neuro-prosthetic system. For epi-retinal prosthetic devices, maximal resolution would ideally involve stimulation of individual RGCs, requiring electrode sizes to match those of their target neurons. However, small electrodes require higher charge densities to provide sufficient power to drive cells above firing threshold. If the charge density is too high, it can damage the tissue or the electrode, rendering the system unusable in the long term. Hence, it is important to estimate safe charge density limits based on electrode and tissue properties, allowing the system to function safely over prolonged periods. An ideal epi-retinal stimulation system would thus include very small capacitive electrodes, located in close proximity to the RGC layer, requiring low amounts of current to depolarise RGCs to threshold.

As charge density is intrinsically related to the effective surface area of the electrode, the geometry of stimulating electrodes strongly affects their charge density limit. As such, materials with large effective surface areas are ideal for efficient stimulation. Materials such as titanium nitride (TiN), iridium oxide (IrOx) and platinum grey are considered as gold standards for neural prosthetics and used respectively by the three main retinal prosthetic projects^5, 6, 7^. Carbon nanotubes (CNTs) have attracted much attention since their emergence in the field of bioengineering^8^ due to their outstanding electrical^9, 10^, chemical and mechanical properties. Their high surface area^11^, remarkable tensile strength^12^, biocompatibility^13^ and high conductivity^14^ make them an alluring candidate material to use in neural prosthetic electrodes^15, 16, 17^.

Neurodegeneration is characterized by strong glial proliferation, and rejection of epi-retinal electrodes by glial cells can potentially widen the gap between electrodes and their target neurons, thus increasing the amount of charge required to elicit action potentials. The glial population of the retina, consisting of microglia (scattered throughout the inner retinal layers), astrocytes (horizontal syncytium at the nerve fibre layer) and Müller cells (spanning the retina transversely) maintain homeostasis^18, 19^, provide immunological protection^20, 21^ and structural support^22, 23^. Macroglia (astrocytes and Müller Cells) are also the source of the inner limiting membrane (ILM), a basement membrane providing physical and electrostatic barrier between the vitreous and the retina^24^. Similar to other basement membranes, the ILM is composed of three layers^25^: the *lamina rara interna* (adjacent to glial cell endfeet), the *lamina densa* and the *lamina rara externa* (contiguous with the vitreous humour). These layers have the typical molecular constitution found in basement membranes with laminins, collagen, and other heparan sulfate proteoglycans^26, 27, 28, 29^. Integration of epi-retinal electrodes into the ILM is essential to reduce the distance between stimulating electrodes and target neurons.

Understanding the interactions between stimulating electrodes and the complex micro-environment at the vitreo-retinal interface is thus an important step for the successful design of efficient epi-retinal prosthetic devices. In this study, we investigated the response of RGCs, macroglial cells and the ILM of dystrophic retinas (here, we used the Crx knockout mouse model of Leber congenital amaurosis^30, 31^) to interfacing with CNT electrodes over up to three days *in vitro*. During this incubation period, we observed a substantial increase in the coupling between RGCs and CNT electrodes, reflected both by electrophysiological and anatomical changes. Signals exhibited a gradual improvement in signal-to-noise ratio and decrease in stimulation thresholds. At the same time, CNT structures became gradually integrated into the ILM, reducing the distance between electrodes and RGCs.

## Results

### Device Fabrication

We fabricated both MWCNT (Multi-walled CNT) MEAs and isolated MWCNT islands (Fig. 1) to investigate respectively the electrophysiological and morphological impact of CNT electrodes interfaced with the inner retina. Active MEAs were used to stimulate cells and to record electrophysiological signals from the RGC layer in intact, live tissue. Passive MEAs were used to study the structural integration of CNT islands initially just loosely attached to a silicon/silicon oxide (Si/SiO_2_) substrate into retinal tissue. The fabrication process for electrically connected CNT MEAs has been described elsewhere^17^, and isolated CNT islands were fabricated using a similar process, but involving fewer steps^32^. Briefly, Ni was patterned on Si/SiO_2_ substrates by photo-lithography, and then used as a catalyst to grow CNTs by Chemical Vapour Deposition (CVD). For active MEAs, the CNT islands had a 30 μm diameter and a 200 μm pitch (30/200). Passive MEAs had two types of islands. 30 μm islands were organised in a central orthogonal lattice and had a 50 μm pitch whilst 100 μm islands were organised in an orthogonal lattice concentric to the 30 μm island lattice (fig. 1b4) with a 200 μm pitch. The charge injection limit of the active CNT electrodes was ~2 mC/cm^2^ ^33^.

**Figure 1.**
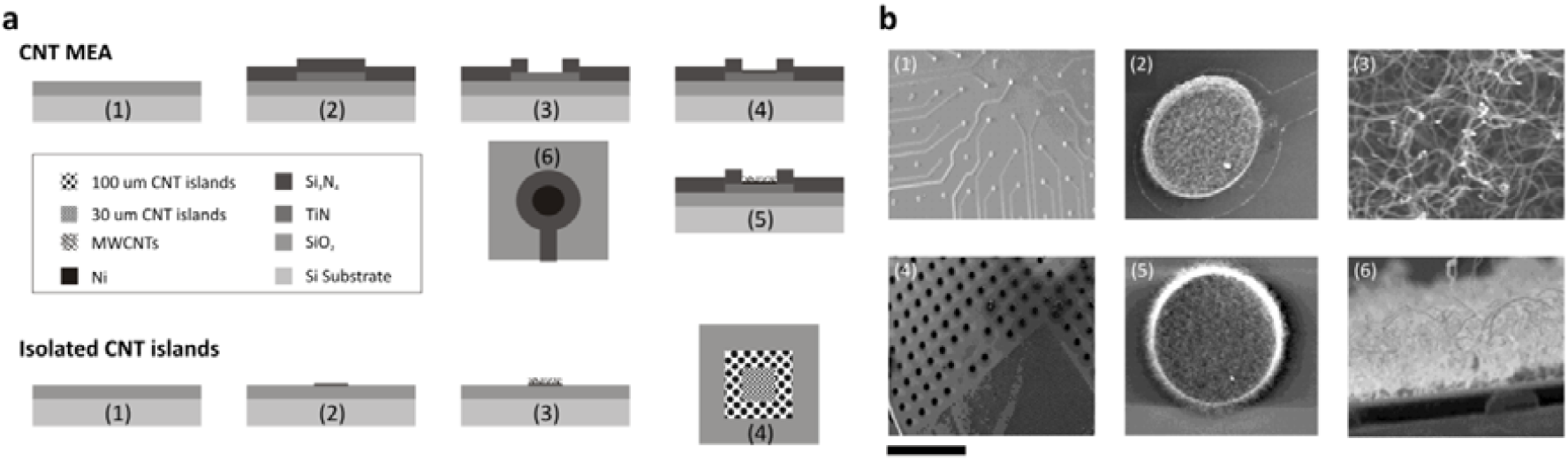
**Fabrication processes for CNT MEAs and loose CNT islands**. **(a)** Manufacturing CNT MEAs (top) begins with a Si/SiO_2_ substrate (1) on which is deposited conductive TiN and passivating Si_3_N_4_ (2). A 30 µm hole is etched into the passivation layer (4) before depositing Ni (4) and growing CNTs by CVD (5), yielding electrically connected CNT electrodes (6; bird’s eye view of single electrode). Isolated detachable CNT islands (bottom) are manufactured by patterning Ni (2) directly onto a Si/SiO_2_ substrate (1) and growing CNTs by CVD (3), yielding concentric lattices of detachable 100 µm and 30 µm CNT islands (4; bird’s eye view of all islands on substrate). **(b)** Scanning electron micrographs at increasing magnification revealing the surface of CNT MEAs **(b1-3)** and loose CNT islands **(b4-6)**, highlighting the large surface area of entangled CNTs. Scale bars are **b1** 500 µm, **b2** 30 µm, **b3** 400 nm, **b4** 1 mm, **b5** 10 µm, **b6** 2 µm.

### Electrophysiological evidence for acute time-dependent increase in coupling of CNT electrodes

To evaluate the performance of CNT electrodes, we compared the electrical behaviour of RGCs coupled to CNT electrodes versus TiN electrodes. Outer retinal degeneration disorders are characterised by the presence of strong spontaneous activity consisting of local field potential (LFP) oscillations and vigorous bursting^32–34^. Fig. 2a,b illustrates these pathological oscillations in the Crx-/- retina^35^ (see next section for more details on the Crx model). We found that these were reduced after 3 days *in vitro* (Fig. 2c), hinting at a breakdown of functionality in the inner plexiform layer under long-term explanted conditions.

**Figure 2.**
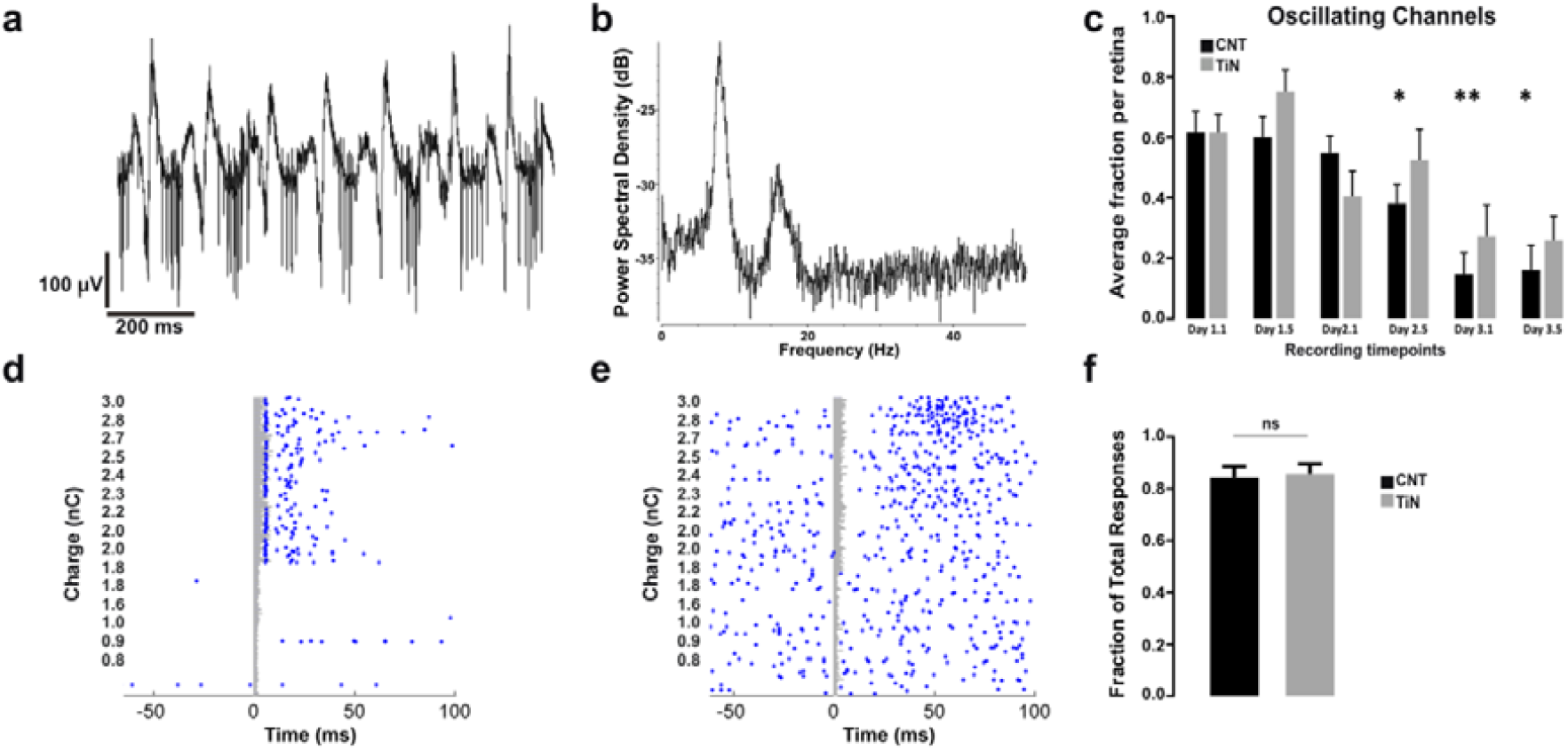
**Spontaneous and evoked retinal activity recorded on CNT and TiN MEAs**. **(a)** Raw extracellular recording from one MEA channel displaying hyperactive bursting and strong LFPs recorded from one electrode on a CNT electrode from the Crx retina. **(b)** Activity power spectrum density plot displayed in **(a)** recorded over the course of 2 minutes, with a peak around 8Hz and its harmonic around 16 Hz. **(c)** Bar graph displaying the proportion of oscillating channels for both types of electrode materials. Wilcoxon matched-pairs signed rank test, comparing fraction of oscillating channels at each time point to that of the first time point. Asterisks indicate significance with p values of 0.0156, 0.0078 and 0.0156 for CNT time points 2.1, 2.5 and 3.5, respectively. N = 6, 6, 6, 6, 4, 4 retinas on TiN MEAs and N = 11, 11, 11, 11, 10, 8 retinas on CNT MEAs for respective time points. **(d-e)** Raster plots displaying examples of direct **(d)** and indirect **(e)** responses recorded on an electrode from a CNT MEA. **f)** Average fraction of direct responses per retina for experiments performed with both TiN and CNT electrodes. Mann Whitney test, p = 0.9609, N = 11 retinas on CNT MEAs and 6 retinas on TiN MEAs.

To differentiate between responses unequivocally elicited by direct stimulation of the spiking RGC from those elicited through stimulation of secondary retinal interneurons making functional connections with the RGCs under investigation, responses were pooled according to latency. Those occurring within 10 ms of electrical stimulation were classified as direct (Fig. 2di) and those occurring after 10 ms were classified as indirect responses (Fig. 2dii). The fraction of direct responses was significantly higher than that of indirect responses for both CNT and TiN electrodes (Fig. 2e) pooled across all recording time points per explant. For the rest of this paper, we concentrate only on these more frequent direct responses.

#### Age-dependent increase in RGC stimulation thresholds in dystrophic retinas

Retinal degeneration can occur naturally (e.g. Royal College of Surgeons rat^34^), be due to genetic alterations or initiated by a physical insult (e.g. retinal detachment or strong light exposure in albino animals). Rodent models of outer retinal dystrophies are widely used^32–35^ to reach a better understanding of the kinetics of retinal degeneration and designing appropriate biological rescue strategies.

Although the kinetics of degeneration differ for each type of retinal dystrophy, retinal outer degenerative disorders follow the same general pattern^32^ (outlined in fig. S1). In this study, we have used the cone rod homeobox (Crx) knockout mouse, a model of Leber congenital amaurosis. Crx is an important gene for the survival of photoreceptors and for the development of their outer segments^30, 31^. Photoreceptors in the Crx-/- mouse are born with atrophied outer segments and they cannot transduce light. They begin to die at postnatal day (P) 30 and the outer nuclear layer progressively shrinks from 12 to 1 layer by P210^35^, resulting in much thinner retinas than normal (fig. S1 and S2). Substantial inner retinal remodelling occurs during that period (fig. S1). The glial scarring associated with degeneration of the Crx-/- retina (fig. 3a) can push stimulating electrodes away from their target RGCs, thus increasing the charge required to bring these neurons to firing threshold.

**Figure 3.**
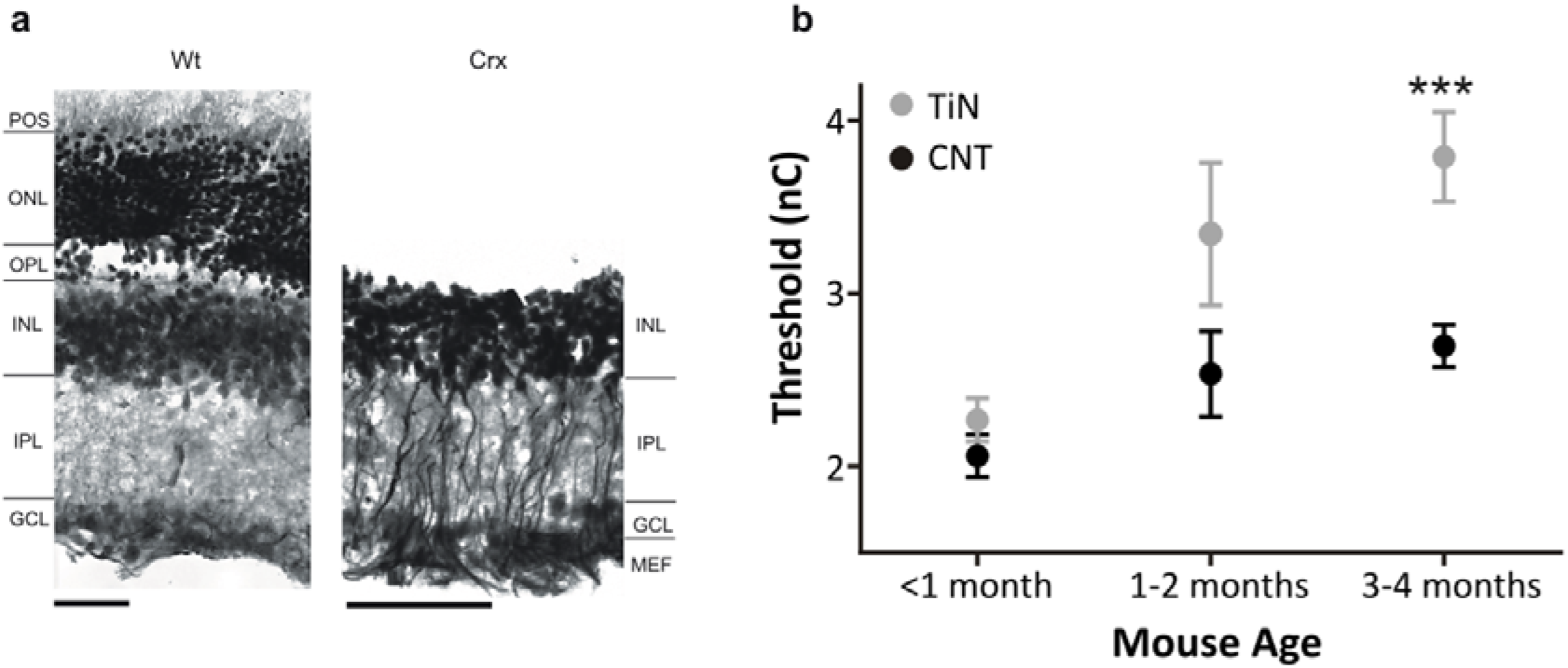
**Age dependent increase in stimulation threshold**. **(a)** Wild type and Crx-/- retinal sections (from 3 months old mice) stained for GFAP with a Nissl counterstain, highlighting the lack of an outer retina and sealing of the inner retina by hypertrophic Müller cells. Scale bars: 50 µm. **(b)** Longitudinal changes in charge thresholds (average ±SEM) for CNT and TiN electrodes in Crx -/- retinas during the first four postnatal months. N (CNT < 1, 1-2 and 3-4 months) = 54, 6, 63. N (TiN < 1, 1-2 and 3-4 months) = 48, 14, 39. ***- p = 0.0002, Mann Whitney test

In order to assess the impact of age-related retinal gliosis on epi-retinal stimulation threshold values, we stimulated RGCs from the Crx-/- retina between birth and 4 months of age. Fig. 3b shows that there is a pronounced increase in threshold charge as degeneration progresses and the neural retina becomes increasingly entangled in glial processes (Fig. 3a). However, the increase in threshold is much less pronounced when using CNT electrodes than when using standard TiN electrodes, with the difference becoming statistically significant at advanced degeneration stages. These results already demonstrate that CNT electrodes offer a more efficient interface for cellular stimulation, presumably because they allow the formation of more intimate contacts between the tissue and the prosthetic device (see below).

#### Spike amplitudes increase with time

The size of action potentials is an indication of the quality of the coupling between the retinal tissue and the recording electrode, as the propagation of such signals through resistive biological tissue is attenuated with increasing distance between the two. We have previously reported that spike waveforms increase in size within the first hour after mounting the retina on a CNT MEA^16^. In this study, we have followed these changes for up to three days *in vitro*, to detect mechanical, electrophysiological and histological changes.

Fig. 4a shows two examples of signal improvement over eight hours on two channels from a CNT array, clearly illustrating that the waveforms become larger with time upon contact with CNT electrodes. Fig. 4b illustrates the cumulative distribution of spontaneous spike amplitudes recorded on a CNT (Fig. 4b) and TiN (Fig. 4c) MEA over three days for one retina. There is a clear shift towards larger amplitudes on Day 2 and Day 3 for the spikes recorded on the CNT MEA, but not for spikes recorded on the TiN MEA. Figure 4c summarizes changes in spike sizes over time for all retinas, clearly showing that CNT, but not TiN electrodes yielded larger signals on Day2 and Day3 than on Day1. Moreover, spikes with amplitudes as large as 400 µV could be seen with CNT electrodes, whereas signals recorded with TiN electrodes never reached such large amplitudes.

**Figure 4.**
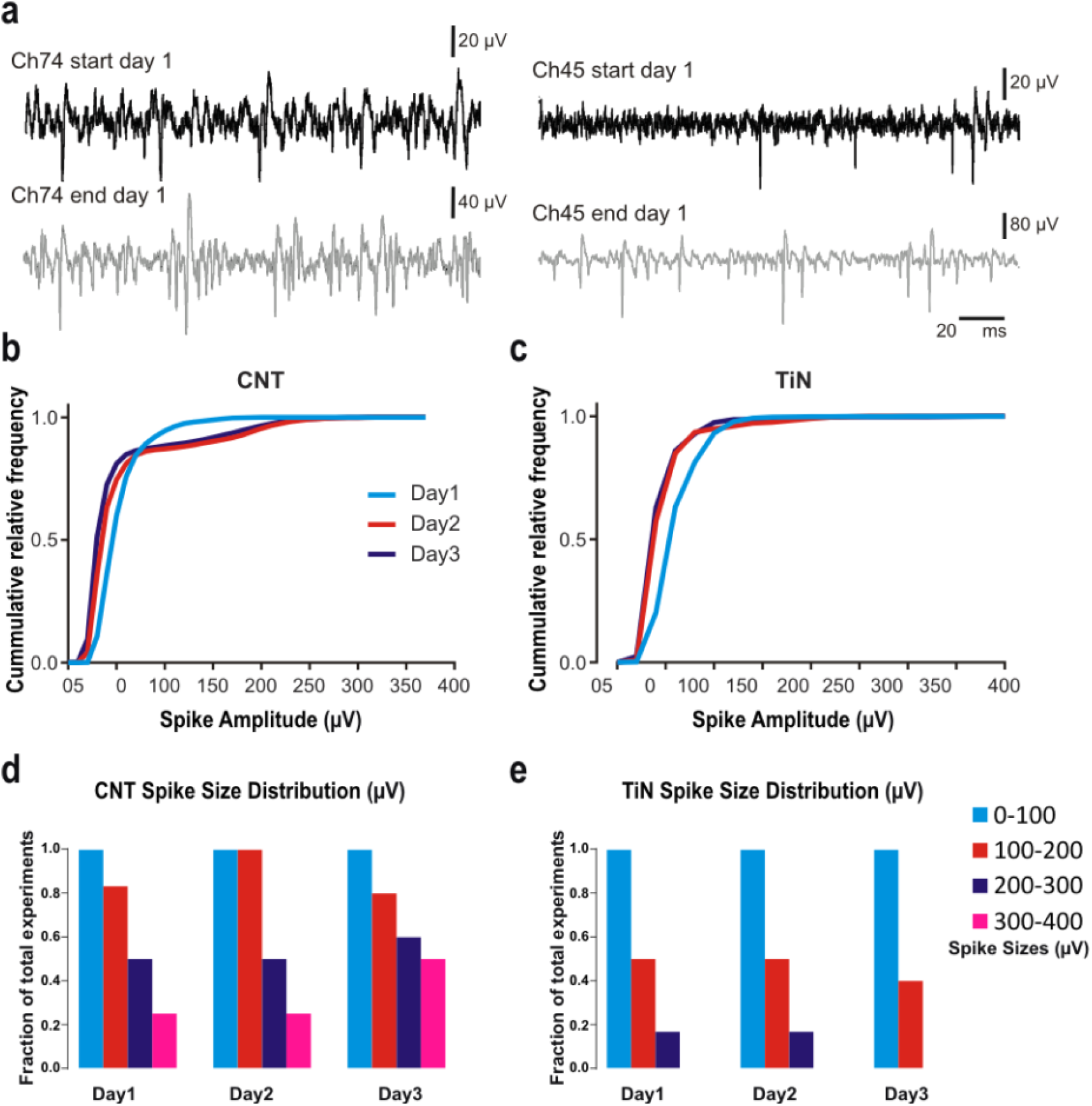
**Acute increase in spike amplitude**. **(a)** Example raw trace displaying spike sizes over the course of eight hours recorded on CNT electrodes. **(b-c)** Relative cumulative distribution of spike amplitudes for a single retina on a CNT **(b)** and TiN **(c)** MEA. **(d-e)** Fraction of retinas which had spike sizes of up to 400 µV on Days 1 to 3 for CNT **(d)** and TiN experiments **(e)**.

#### Stimulation thresholds decrease with time

Seventeen retinas were experimented on over a period of three days (6 on TiN and 11 on CNT MEAs), stimulating the same four individual electrodes (for each retina) on each day, generating a theoretical total of 204 stimulation runs. However, some experiments did not successfully yield data for three consecutive days (e.g. artefacts too strong to remove, tissue deterioration on Day3, human error during experimental procedure), leading to a total of 168 stimulation runs (109 for CNT and 59 for TiN electrode experiments).

Stimuli with a cathodic initial phase and an asymmetric shape consisting of a reversal phase twice as long and with half the current amplitude of the initial phase were significantly more efficient at activating RGCs (lower threshold charge, Fig. S5) than other stimulus shapes both for CNT and TiN electrodes.

Figure 5a illustrates an example of direct responses in a single RGC, requiring an increasingly lower amount of charge for activation over time, whilst concurrently spikes increase in amplitude (Fig. 5b). The fraction of responding RGCs with thresholds < 2nC out of all spiking RGCs per retina increased significantly on Day2 for CNT electrodes but not for TiN electrodes (Fig. 5c). A 2 nC threshold was selected here because some retinas required larger amount of charge to elicit responses than others whilst all retinas were stimulated by at least 2 nC. The average threshold value yielded by CNT electrodes was also significantly lower on each day compared to values obtained with TiN electrodes (Fig. 5d). Finally, when stimulating with CNT electrodes, the average direct threshold values became gradually and significantly lower from Day 1 to Day 3 (Fig. 5e) with no significant changes over time for TiN electrodes. For both electrode materials, fewer cells participated in the responses on Day3, presumably because of deteriorating health of the tissue. These results indicate that for CNT electrodes, gradually less current is required with time to directly depolarise RGCs past their firing threshold, suggesting that the retina becomes increasingly more intimately coupled to the CNTs. This is further confirmed by the higher recruitment rate on Day 2 for CNT electrodes.

**Figure 5.**
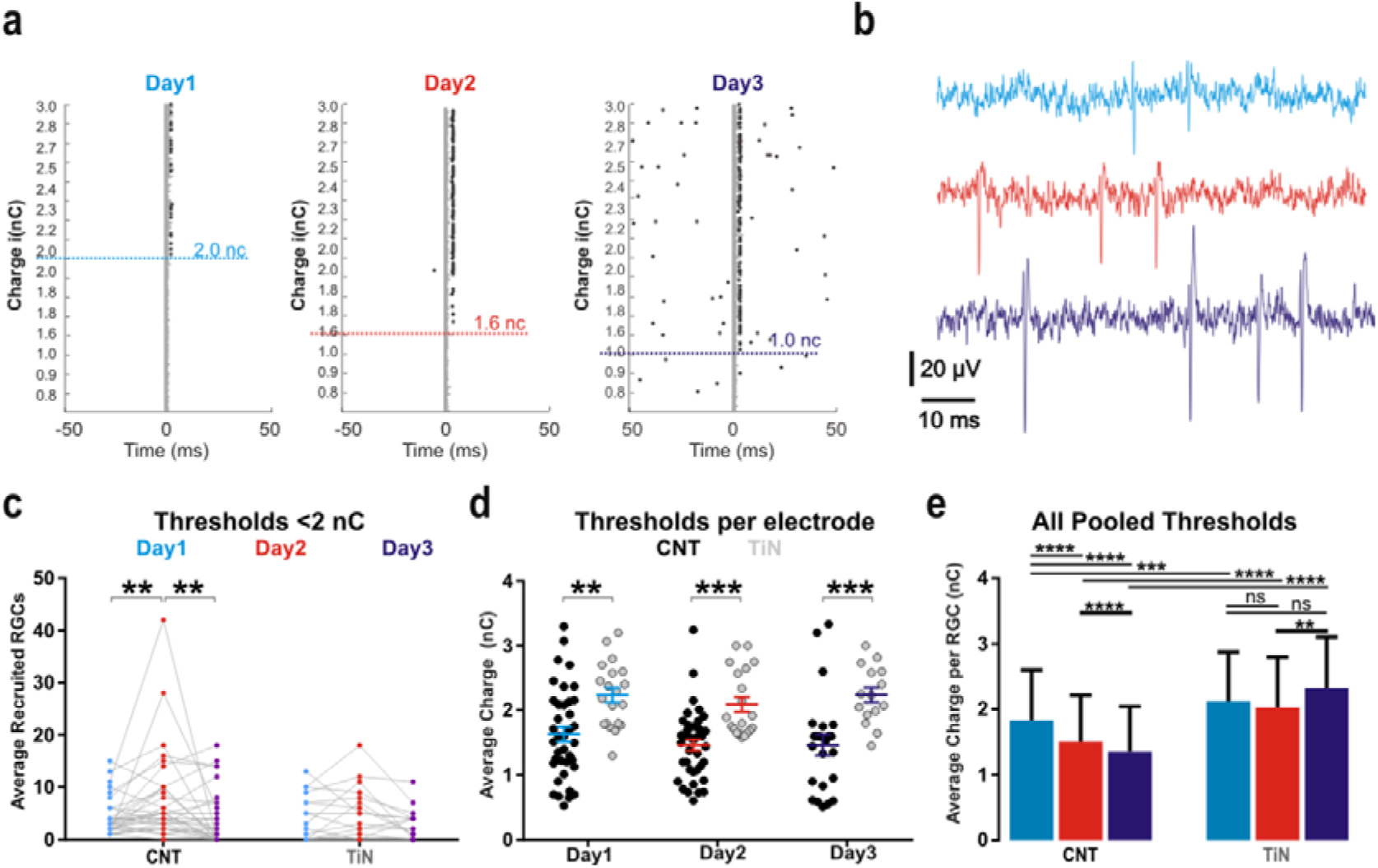
**Changes in stimulation thresholds, spike amplitudes and cellular recruitment over the course of 3 days *in vitro***. **(a)** Raster plots of direct electrical stimulation of an RGC on a CNT MEA, displaying a gradual decrease in charge threshold over time. Epochs are organised by increasing amount of stimulation charge value along the ordinate axes. These are not linear or unique as they represent the product of multiple parameters for stimulus current and single-phase duration (see Methods). **(b)** Raw signal displaying spike waveforms on Days 1-3 from the unit in **(a)**. **(c)** Average number of recruited RGCs with thresholds below 2 nC per electrode on Days 1-3 for both CNT and TiN MEAs. Asterisks indicate statistical significance (Bonferroni’s multiple comparisons test), with p = 0.0081 and p = 0.0022 for CNT Day1-Day2 and Day2-Day3, respectively. N = 34. **(d)** Average threshold values per electrode recorded on Days1-3 for both types of MEAs. Asterisks indicate statistical significance (Sidak’s multiple comparisons test), with p = 0.0013, p = 0.0008 and p = 0.0006 for Day1 Day2 and Day3, respectively. For CNT electrodes, N = 38, N = 39 and N = 23 for Day1, Day2 and Day3, respectively. For TiN electrodes, N = 20, N = 20 and N = 15 for Day1, Day2 and Day3, respectively. **(e)** Average (±SD) threshold (nC) across all RGCs recorded in this study. Asterisks indicate statistical significance (Dunn’s multiple comparisons test), with ***p < 0.0001 and **p = 0.0059. N = 241 (CNT Day1), N = 389 (CNT Day2), N = 155 (CNT Day3), N = 230 (TiN Day1), N = 287 (TiN Day 2) and N = 148 (TiN Day3).

### Anatomical evidence for time-dependent increase in coupling of CNT electrodes

The morphological and anatomical effect of CNT assemblies on the inner retina were investigated with immunohistochemical (IHC) labelling and electron microscopy performed on retinal explants which had been interfaced with CNT constructs for 4, 12, 24 and 72 hours (see Methods and fig. S6). CNT islands initially loosely attached to their Si/SiO_2_ substrate could eventually be stripped off their substrate as they became gradually embedded in the retina. Semi-thin sections of retinal bio-hybrids were used to quantify the physical coupling between the retinas and CNT islands (we were not able to gather any transmission electron microscopy (TEM) data for 30 μm islands as none were found during the semi-thin sectioning process). We used two parameters to quantify coupling, one is the average number of islands adhering to each retinal explant (fig. 6c) and the other is the distance between the external surface of each island and the retinal ILM (fig. 6d). Similar measurements were taken on all islands adhering to retinas for each particular time point. 100 μm CNT islands gradually integrated into the ILM over 48 hours (fig. 6), evidenced by a decrease in the distance between the CNT islands and the tissue (Fig. 6a, b and d) and by an increase in the number of islands adhering to the retina (Fig. 6c). As there were only few 100 μm adhering islands at shorter time points, we could not perform reliable statistical analysis on that particular dataset.

**Figure 6.**
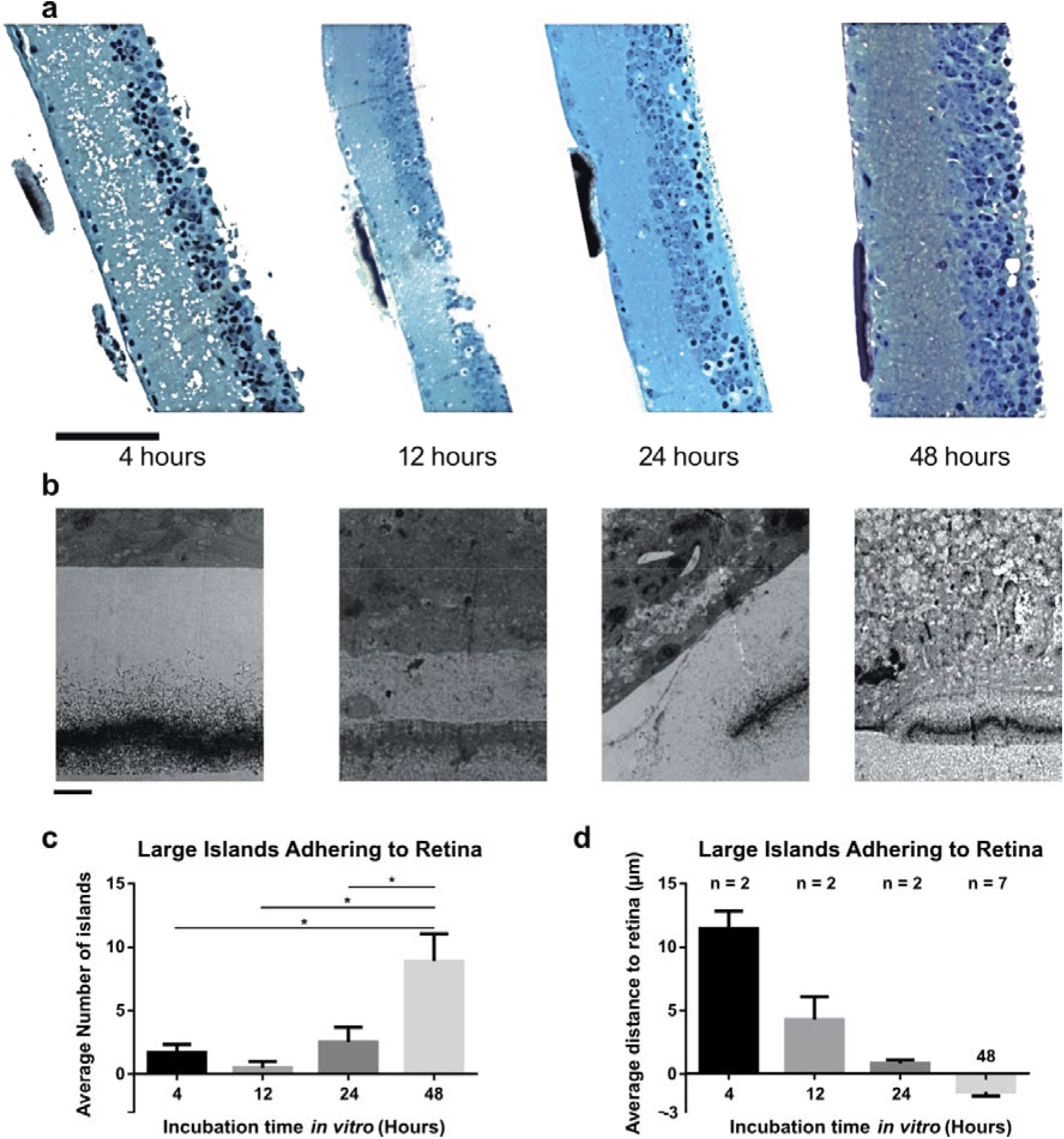
**Integration of large CNT islands into retinal explants over 48 hours**. **(a)** Semi-thin transverse sections stained with Toluidine blue demonstrate the progressive integration of CNT islands into the inner retina over four time points (4, 12, 24 and 48 hours). Scale bar is 100 µm. **(b)** TEM micrographs showing the ultrastructural details of the integration described in **(a)**. Scale bars are 2.54 µm (4 hours), 5 µm (12 hours), 3.36 µm (24 hours) and 8.4 µm (48 hours). (c) Bar graphs demonstrating the progressively higher number of large CNT islands adhering to retinas with time (average ±SEM). Asterisks indicate statistical significance (Mann-Whitney test), with p = 0.0318, p = 0.0364, p = 0.0434 for 48h vs 4, 12 and 24h, respectively. **(d)** Bar graphs demonstrating the progressively shorter distance between large CNT islands and retinal tissue with time (average ±SEM). The negative values highlight the fact that portions of the islands are physically engulfed in the ILM. It was not possible to perform statistical tests because too few CNT islands adhered to the retina at short incubation times.

TEM performed at each of these four time points revealed a more detailed picture of the integration of CNT islands within the ILM (Fig. 6b). After 4h, there was no direct contact between individual CNTs and the ILM (fig. 6b, left panel). Indeed, the closest CNT we found to the ILM was approximately 2 µm away (fig. S7a, blue arrowhead). We also observed some strands of biological material (fig. S7a, green arrowhead) between the CNTs and ILM, suggesting that some remnants of the vitreous humour (probably collagen fibrils) were buffering the zone between the electrode and the retina. After 12h, we found evidence of thickening of the vitreous between the retina and the CNT island. Figure S8 illustrates this process, with portions of the vitreous closer to the retina appearing denser (fig. S8f) than those further away (fig. S8e). Indeed, there are fewer and smaller gaps (white portions of the micrograph which have not been stained with osmium tetroxide, uranyl acetate or lead citrate) in the vitreal matrix in panel f than in panel e. After 24 hours in contact with CNT islands, the retina changed shape to accommodate the island (fig. S9a). We also observed evidence for the formation of an accessory limiting membrane (fig. 6b and s9b) which grasped the CNT island from the edges and on occasions, we could see individual CNTs penetrating the ILM.

After 48 hours, islands were completely integrated into the ILM (fig. 6b), with collagen fibrils from the lamina rara externa anchoring individual CNTs (fig. s10) and the accessory limiting membrane described above extending for tens of microns. Fig. 7 shows a transverse profile of how the retinal tissue interacts with a CNT island at increasingly higher magnification. Panel a shows a light micrograph of a toluidine blue semi-thin section, already clearly indicating that the CNT island has become integrated in the retinal tissue, which is more conspicuous on the lower power TEM in panel b. Panel c is a magnification of the edge of the CNT island, where individual CNTs can be seen enveloped in a homogenous matrix. This matrix can be seen extending for tens of microns along the ILM (d), becoming progressively thinner as the distance from the CNT island edge increases. We also observed multiple instances of CNTs penetrating the retinal tissue (fig. 7e).

**Figure 7.**
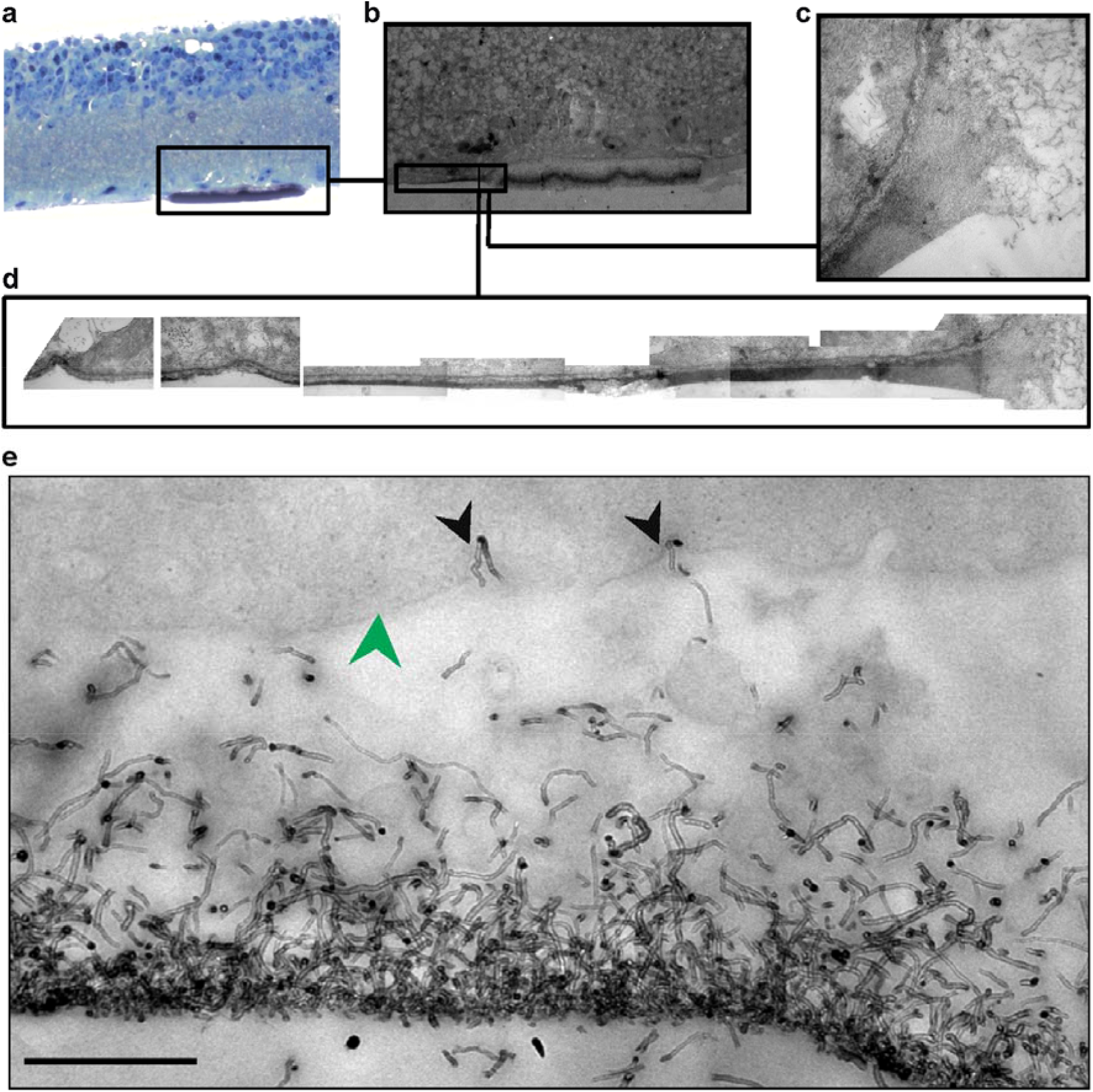
**Homogenous matrix grappling edge of CNT island after 48 hours *in vitro* extends along the retina for over 24 μm**. **(a)** semi-thin transverse section stained with Toluidine blue. **(b)** TEM micrograph of the same retina with the CNT island grasped by a long homogenous matrix (box). **(c)** Higher magnification micrograph showing the homogenous matrix grabbing the edge of the CNT island. **(d)** digitally-stitched collage of 10 TEM micrographs which follow the matrix to an invagination in the ILM 23 μm away. **(e)** Individual CNTs (black arrowheads) penetrating the retinal ILM (green arrowhead). Scale bar is 50 μm in **(a)**, 60 μm in **(b)**, 3 μm in **(c)**, 4.5 μm in **(d)** and 500 nm in **(e)**.

Scanning electron micrographs (SEM) of explants incubated over 48 hours revealed both 50 μm (fig. 8a) and 30 μm (fig. 8b) CNT islands incorporated into the ILM. The panel in the top-left quadrant of fig. 8a displays an explant with ten 100 μm islands adhering to its surface, covered by the ILM which has a characteristically smooth appearance in SEM (Fig. S3). The two middle panels focus on one of the islands at higher magnification, revealing fibrous bundles (blue arrows) clutching the side of the island. These fibre bundles can reach tens of microns in length, perhaps reflecting the long matrix we describe in fig. 7(a-d), although they may also reflect activated Müller cells (see fig. 9c). The panel in the bottom-right quadrant focuses on part of the island periphery, which appears to be covered in an accessory limiting membrane with a texture somehow similar to the ILM (green arrow), with gaps exposing the CNTs underneath (yellow arrows). The bottom left panel focuses on a CNT island which has been fractured following critical point drying (Methods), allowing us to visualize biological bundles (as described above) infiltrating the CNT island throughout.

**Figure 8.**
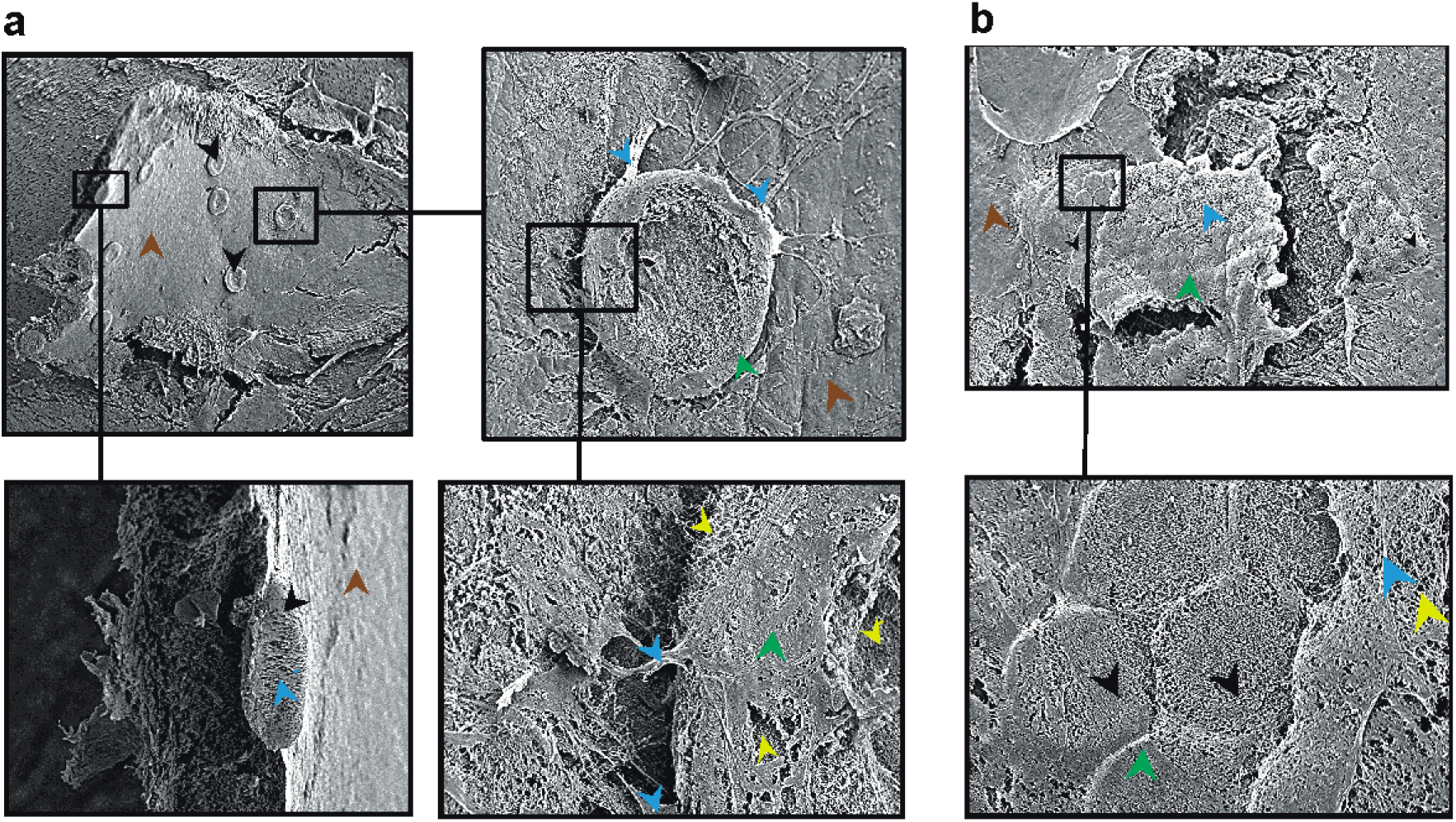
**SEM images of the ILM incorporating large CNT islands**. **(a)** Retinal explant with 10 large CNT islands (black arrowheads) adhering to its ILM (brown arrowheads). Scale bars are 847.6 µm (top-left quadrant), 100 µm (top-right quadrant), 125.35 µm (bottom-left quadrant), and 23.42 µm (bottom-right quadrant). **(b)** Retinal explant with over 50 small CNT islands (black arrowheads) embedded into an accessory limiting membrane (green arrowhead) continuous with the ILM (brown arrowhead). Scale bars are 200 µm (top panel) and 27.38 µm (bottom panel).

Fig. 8b displays a portion of the retina covered by more than fifty 30 μm CNT islands, which have formed in a hexagonal lattice as they are incorporated into the retina. When viewed at higher magnification (bottom right panel), some islands are not completely covered, allowing visualization of individual CNTs (yellow arrowhead) below ILM fibres (blue arrowheads) in the process of weaving the accessory limiting membrane.

Immunohistochemical labelling of wholemount retina-CNT biohybrids with Glial Fibrillary Acidic Protein (GFAP) allowed to estimate the impact CNT assemblies have on retinal glial populations. In response to an invasive threat, glial cells are known to activate, a process characterised by the upregulation of GFAP^36, 37^ and followed by various cellular responses which can be detrimental to, and even reject an implanted device^38^. In the dystrophic retina, Müller glia are in a state of activation and heavily involved in the remodelling process associated with degeneration^39, 40^ (fig. S1 and S2). Tears in the ILM lead to reactive gliosis, emphasizing the requirement for gentle interfacing between epi-retinal electrodes and ILM. Fig. 9a displays a positive control for gliosis, with fluorescent double labelling for laminin (one of the main components of the ILM) and GFAP, highlighting an increase in GFAP levels coinciding with the areas showing tears in the laminin layer. The average number of 100 μm CNT islands initialising reactive gliosis per retina was significantly larger than for 30 μm islands (fig. 9e). 100 μm CNT islands induced both astrocytic (fig. 9b) and Müller gliosis (fig. 9c) whilst 30 μm islands were rarely seen to promote gliosis (fig. 9d).

**Figure 9.**
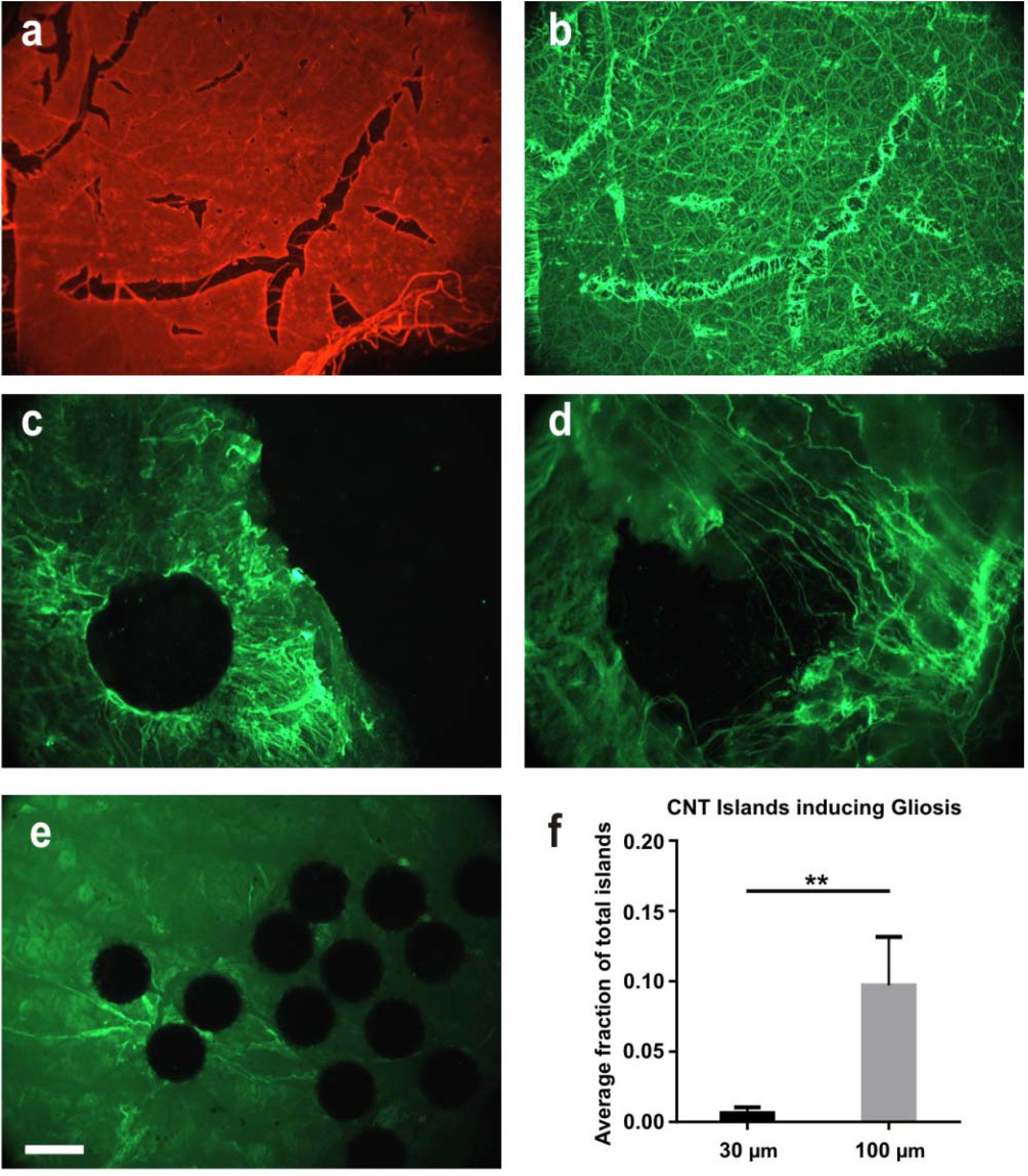
**Immunhistochemical analyses of CNT-retinal interactions**. **(a-b)** Micrographs of control retinal wholemounts stained with laminin **(a)** and GFAP **(b)**. **(c-d)** Higher magnification micrographs of retinal wholemounts interfaced with 100 µm CNT islands and stained for GFAP, displaying activation of both astrocytes **(c)** and Muller cells **(d)**. e) micrographs of retinal wholemounts interfaced with 30 µm CNT islands and stained for GFAP. **(f)** Bar graph displaying the average fraction of islands inducing reactive gliosis. Asterisks indicate significance (Mann-Whitney test), with p = 0.0064, N = 7 explants. Scale bar is 75 µm for **(a-b)**, 30 µm for **(c-d)** and 12 µm for **(e)**.

## Discussion

In this study, we have maintained dystrophic retinas for up to three days *in vitro* on CNT and TiN electrodes, demonstrating for the first time that CNTs become gradually incorporated within the ILM without promoting reactive glial responses, resulting in gradual decrease in stimulation thresholds and increase in the number of RGCs reaching firing threshold. Although previous studies have already demonstrated that CNTs offer great potential for epi-retinal prosthetic devices^16, 41, 42^, here we provide the first comprehensive evaluation of how CNTs interact with RGCs in the diseased retina, comprising ultrastructural, immunohistochemical and electrophysiological evidence, demonstrating a clear correlation between temporal changes in structure and function underlying the ability of CNT electrodes to stimulate RGCs and how it improves with time. Collectively, our data indicate that CNTs are indeed a promising electrode material for retinal prosthetics, with superior biocompatibility and electrical properties.

The ability to safely stimulate RGCs requires implanted electrodes to deliver supra-threshold charge over prolonged durations without causing damage to the tissue or electrode material. Although this goal has been generally achieved, one of the technological hurdles remaining is the size of the electrodes, with smaller electrodes leading to higher spatial resolution. But electrodes with a smaller surface also require higher stimulation charge density to reach supra-threshold levels. Not only are electrode materials intrinsically limited by the density of charge they can store at their surface, but high charge densities can lead to electrode and tissue damage^4, 43, 47^. Lower charge injection during electrical stimulation would lead to more energy efficient prostheses, thus increasing the potential number of powered channels as well as battery life and computational power. Lower charge densities can be achieved by (1) using “ideal” stimulating electrode materials with a larger surface area; (2) using optimal stimulation protocols requiring less charge to depolarise target cells; and (3) reducing the distance between electrode and target neurons. These points will be discussed below with reference to the data presented in various studies, including the present one (summarized in Table 1).

**Table 1.** **Comparison of performance between different studies using electrical retinal stimulation**. Thresholds are highest *in vivo* and for sub-retinal electrodes. Underlined numbers were derived from data provided in the referenced papers. Electrode spacing, h: horizontal, d: diagonal.

| Reference | Experimental setup | Species | Electrode placement | Electrode material | Electrode diameter/spacing ($\mu\text{m}$) | Pulse Width ($\mu\text{s}$) | Threshold charge density ($\text{mC}/\text{cm}^2$) |
| --- | --- | --- | --- | --- | --- | --- | --- |
| <sup>51</sup> | *In vitro* | Rabbit | Epi-retinal Healthy | Pt-Ir | <u>3.56</u>/Na | 100 | <u>5-32.14</u> |
| <sup>7</sup> | *In vivo* | Human | Epi-retinal Dystrophic | Pt Grey | 200/502.5d | 450 | 0.29 |
| **This study** | *In vitro* | Crx-/- mouse | Epi-Retinal Dystrophic | TiN | 30/200 | 10-100 | <u>0.27</u> |
| **This study** | *In vitro* | Crx-/- mouse | Epi-Retinal Dystrophic | CNT | 30/200 | 10-100 | <u>0.22</u> |
| <sup>56</sup> | *In vivo* | Human | Sub-retinal Dystrophic | TiN | 112.83/280h<br>396d | 2000-3000 | <u>0.2-0.6</u> |
| <sup>57</sup> | *In vitro* | Mouse rd1 | Epi-retinal Dystrophic | TiN | 30/200 | 100-1000 | 0.235-0.353 |
| <sup>58</sup> | *In vitro* | Rat/RCS rat | Sub-retinal Dystrophic | IrO<sub>2</sub> | 60, 40,<br>20/285, 145,<br>75 | 4000 | <u>0.26-0.32</u> |
| <sup>54</sup> | *In vitro* | Rat | Epi-retinal Healthy | Pt | 7-16/60 | 50/100 | 0.058 |
| <sup>59</sup> | *In vivo* | Human | Epi-retinal Healthy | Pt | 500 and<br>250/800 | 1000 | 0.05-0.57 |
| <sup>52</sup> | *In vitro* | Rhesus monkey | Epi-retinal Healthy | Pt | 9-15/60 | 50/100 | 0.05 |
| <sup>54</sup> | *In vitro* | Rat P23H | Epi-retinal Dystrophic | Pt | 7-16/60 | 50/100 | 0.049 |
| <sup>41</sup> | *In vitro* | Rat | Epi-Retinal Healthy | PEDOT-CNT | 30/200 | 200 | <u>0.0032-0.0159</u> |
| <sup>41</sup> | *In vitro* | Rat | Epi-Retinal Healthy | TiN | 30/200 | 200 | <u>0.0159-0.0318</u> |

### Ideal stimulating electrode materials

The reversible charge storage capacity depends on the electrode material, its size and shape, composition of the electrolyte, and the parameters of the stimulation waveform. As such, materials with large effective surface areas make ideal stimulating electrodes, and CNTs offer a number of substantial advantages over their metal-based counterparts. Mainly, their fractal-like geometry results in a large surface area, significantly increasing the charge injection capacity and lowering the electrode impedance value^44^. Conventional MEAs can be easily coated with solutions of single wall CNTs and used for the recording of action potentials in RGCs^42^. A recent study demonstrated the functionalization of PEDOT with single wall CNTs for retinal stimulation^41^, achieving thresholds as low as 1 nC for PEDOT-CNT electrodes and 5 nC for TiN electrodes. The coatings described in those two studies are ideal for *in vitro* proof of concept experiments, but may delaminate over time, making them unsuitable for long-term clinical applications. The CNTs constituting our electrodes are grown by CVD directly into the porous TiN base, making them highly stable and unable to delaminate as a result of sustained electrical stimulation.

### Ideal stimulation protocols

Epi-retinal stimulation of second order retinal neurons leading to synaptically driven responses has recently generated some interest^41, 45^ with the optimisation of stimulation protocols based on longer pulses^46^. However, because of anatomical evidence pointing to remodelling of the inner retina during degeneration^39, 40^ combined with the fact that long stimulation pulses lead to irreversible faradaic reactions^47^, we chose to restrict our stimulation parameters to shorter pulses (<100 µs), targeting RGCs directly. Although CNT electrodes proved effective over short stimulation timescales, stimulus pulses longer than 500 µs led to amplifier saturation, making it impossible to calculate the chronaxie and establish a rheobase.

Our stimuli were always charge balanced to avoid irreversible faradaic reactions at the electrode-electrolyte bilayer. However, the shape and polarity of each stimulus was delivered in a pseudo-random order. Our experiments yielded consistently lower thresholds when stimulating with (1) a cathodic first phase as well as (2) a slow reversal potential (second phase twice the duration and half the amplitude of the initial phase). A small proportion of our responses (<1%) consisted of axonal stimulation identified by their increase in amplitude with increased charge, a typical characteristic of electrically evoked fibre bundle volleys.

Modelling studies demonstrate that RGC somata have lower thresholds than their axons^48^ owing to the denser distribution of voltage gated sodium channels at the initial segment^49, 50^. This is supported by *in vitro* stimulation of the rabbit retina^51^ with a 10 μm^2^ Pt-Ir extracellular electrode which yielded somatic RGC thresholds 50% lower than axonal thresholds. *In vitro* stimulation of rhesus monkey retinas with closely spaced Pt-plated electrodes (60 μm pitch, 9-15 μm diameter) presents evidence that the area most sensitive to epi-retinal stimulation is close to the soma and the proximal portion of the axon^52^. Patients implanted with the Argus I and II epi-retinal prostheses (Second Sight Medical Products, Inc, Sylmar, CA) report streak-like visual percepts rather than more commonly reported punctate-shaped phosphenes, suggesting direct stimulation of axonal bundles. This is to be expected considering that this study used very large electrodes (200 μm diameter) located far from the retina (~180 μm), necessitating over 90 nC to reach RGC stimulation thresholds^7^.

The use of much smaller CNT electrodes integrated to the patient’s ILM would considerably reduce the amount of charge required to elicit phosphenes, and thus may contribute to reducing the probability of activating axonal responses.

### Reducing RGC-Electrode distance

The present study has demonstrated the ability of CNT constructs to progressively integrate the inner retina whilst improving biophysical interactions between RGCs and electrodes without promoting a glial response. Indeed, gliosis was only experienced with 50 μm but not 30 μm CNT islands, the later having a comparable size to the CNT electrode contacts in our electrophysiological experiments. The use of detachable CNT assemblies interfaced with the retina to estimate GFAP levels proved challenging. Indeed, as they are completely opaque, CNT islands prevented the visualisation of glial processes between CNTs and the RGC layer. However, we were able to visualize gliosis occurring at the edges of the CNT islands as well as “behind” them, i.e. on the vitreal surface of the retina, as glial cells appear to engulf the CNT islands (which is also suggested by our SEM pictures). Labelling retinal explants post recording was unfortunately not feasible because the process of removing the delicate retinal explants (these retinas are extremely thin, being devoid of photoreceptors) from the CNT contacts of the MEA resulted in severe lacerations in the retinal tissue, as CNTs were tightly bound to the ILM by the CNT “Velcro” effect demonstrated here through electron microscopy.

It is this Velcro effect, noticeable even just shortly after mounting the retina on CNT MEAs^16^ that prompted us to investigate longer term effects. The ultrastructural results we present in this study demonstrate that four hours after mounting the retina onto CNT arrays, there is no direct contact between CNTs and the ILM. This suggests that the most viscous portion of the vitreous is still firmly implanted into the ILM and may necessitate several hours of aCSF perfusion to dissolve away. One approach to improve implantation of a retinal prosthetic device would be to digest portions of the vitreous and ILM prior to implantation with glycosidic enzymes^53^.

The strong adhesion that develops with time between the CNT assemblies and the vitreo-retinal interface can solve one of the fundamental problems with epi-retinal prosthetics, which is how to fix the prosthetic device to the retinal tissue. Currently, fixing arrays of stimulating electrodes to the inner retina requires the use of retinal tacks, which introduce a large gap between the electrodes and the tissue, resulting in very high stimulation (perceptual) thresholds, as high as 90 nC^7^ (0.28 mC/cm^2^), a value several orders of magnitude higher than thresholds obtained in our study. Table 1 displays a list of studies investigating different types of electrodes for epi-retinal stimulation. The lowest values, with thresholds below 0.1 mC/cm^2^ are recorded in healthy retinas (except for Sekirnjac^54^ et al., who were able to record evoked responses on the stimulating electrode as well by using a stimulation blanking system which we did not have for our study). This provides additional evidence as to the impact of reactive gliosis (discussed below) on epi-retinal stimulation of RGCs.

Residing directly between epi-retinal electrodes and their target cells, astrocytes represent a barrier to efficient direct RGC stimulation because a hypertrophic glial response to the insertion of a foreign body would widen the gap, thus increasing activation thresholds. Although quiescent astrocytes provide a neuroprotective environment for RGCs under normal conditions, they have been shown to promote RGC injury following activation^55^. In the retina, the target neurons are protected by the ILM and surrounded by Müller cells, astrocytes and blood vessels. An ideal epi-retinal electrode would penetrate the ILM, bypassing the glial cells without provoking unspecific reactive gliosis and promote intimate contact with RGCs. This is exactly what our CNT electrodes appear to achieve, allowing us to conclude, based on our electrophysiological, structural and immunological data, that CNT electrodes offer a promising alternative for the design of new generation epi-retinal prosthetics.

## Methods

### Retinal isolation

Experimental procedures were approved by the UK Home Office, Animals (Scientific procedures) Act 1986. Animals were sacrificed by cervical dislocation followed by immediate enucleation. Isolation of the retina was performed in carboxygenated (95% CO_2_/5%CO_2_) artificial cerebro-spinal fluid (aCSF) containing (in mM): 118 NaCl, 25 NaHCO_3_, 1 NaH_2_PO_4_, 3 KCl, 1 MgCl_2_, 2 CaCl_2_, 10 C_6_H_12_O_6_, 0.5 L-Glutamine (Sigma Aldrich, UK). First, the cornea was pierced with a hypodermic needle (23G gauge, BD Microlance 3, USA). Then, a pair of Vannas scissors (WPI, USA) was used to shear through the cornea, along the ora serrata before removing the lens. The sclera was then peeled off, using two pairs of sharp forceps, exposing the retina. The vitreous humour was teased off using those forceps, taking great care not to pierce the retina or remove the ILM.

### Electrophysiological setup

Isolated retinas were mounted onto MEAs with the RGC layer facing down onto the electrodes and the optic disc outside the MEA active area. To improve coupling between the tissue and the electrodes, a polyester membrane filter (5μm pores) held the retina in place whilst being weighed down by two stainless steel anchors bearing a framework of parallel glass capillaries. Thirteen retinas were mounted onto CNT MEAs and six onto TiN MEAs (MultiChannel Systems, Reutlingen, Germany).

To maintain physiological conditions, the tissue was perfused with oxygenated aCSF at 1 ml/min over the course of up to 72 hours using a peristaltic pump (SCI400, Watson Marlow, UK). During the recording of electrophysiological activity, retinal explants were maintained at 32°C using a temperature controller (TC02, MultiChannel Systems, Reutlingen, Germany) regulating a metallic plate below the MEA (MEA1060 INV, MultiChannel Systems, Reutlingen, Germany) and an inline heater for the inflow of aCSF (Ph01, MultiChannel Systems, Reutlingen, Germany). The experimental setup is outlined in Fig. S4. To prolong the viability of the tissue, retinas were kept at room temperature overnight^60, 61^.

Although Crx-/- mice do not develop photoreceptor-mediated vision, melanopsin-induced phototransduction occurs in some RGCs in the presence of blue light^62^. To avoid activation of these cells, all experiments were carried out in complete darkness.

Electrophysiological signals were recorded using 60-channel MEAs interfaced with a computer running the proprietary software MC_Rack (MultiChannel Systems, Reutlingen, Germany) using an MEA1060INV (Multi Channel Systems, Reutlingen, Germany) amplifier via a PCI MCS card.

Recordings were acquired at 25 kHz sampling frequency. Spontaneous activity was recorded for 10 minutes before and after each stimulation run, amounting to 15 stimulation runs and 15 spontaneous activity files per retina. Electrical stimulation of retinal tissue was achieved via individual electrodes on the MEA connected to an STG2002 stimulator (Multi Channel Systems, Reutlingen, Germany) controlled by a computer running the proprietary software MC_Stim (Multi Channel Systems, Reutlingen, Germany). The stimulator was also connected to the recording equipment’s PCI acquisition card for synchronised recordings. Acquisition of electrophysiological signals during stimulation runs included 100 ms before and 200 ms after the stimulus.

Retinas were electrically stimulated by rectangular charge-balanced biphasic pulses with a 30 µs inter-phase interval (Fig. S5a). The charge delivered by each pulse was modulated by altering either the current intensity (10-100 µA per phase) or phase duration (10-100 µs per phase).

### Analysis of electrophysiological data

MCD files were imported into Matlab (The MathWorks, USA) using the FIND toolbox^63^. The artefact introduced by applying electrical stimulation often saturates the amplifier for a few ms, followed by a large voltage deflection. This artefact was removed by local curve fitting using the SALPA algorithm^64^. Spikes were automatically extracted, then sorted by supra-paramagnetic clustering and wavelet analysis using Wave_clus^65^.

Epochs containing sorted spikes for each electrode were mapped to the different stimulus conditions and organised as raster plots with the ordinates divided into the different charge injected values in ascending order. Consequently, the results for a single stimulation run consisted of a set of six (the number of different stimulating combinations) raster plots for each spike cluster on each recording electrode.

Evoked responses were identified on the raster plots by their synchronicity in repetitive trials of a given condition and by their disappearance below a given amount of injected charge), determining the firing threshold for eliciting spikes (fig. 2d). Spike sizes were measured using the peak-to-peak amplitude of each extracted waveform. Statistical analyses and graph plotting were carried out using Sigma Plot (Systat Software Inc., San Jose, California, USA) and Excel (Microsoft, USA) respectively. Statistical significance was measured using non-parametric analyses with a significance limit of p ≤ 0.05.

### Viable Retina-CNT Biohybrids

In order to investigate the morphological coupling between CNT constructs and the inner retina, retinas were incubated laid on passive CNT discs loosely attached to their Si/SiO_2_ substrate (fig. S6). Incubation chambers consisted of a 100 mm diameter Petri dish with multiple (4 and above) plastic rings (6mm height, 10mm diameter) affixed to the bottom with polydimethylsiloxane (PDMS), a biocompatible silicone adhesive^66^. Passive CNT devices were glued to the bottom of Petri dishes. In order to render the CNT islands hydrophilic, incubation chambers were initially ionised with Nitrogen plasma for 10 minutes and submerged in de-ionised H_2_O for 24 hours prior to retinal incubation in aCSF. Isolated retinas were placed, RGC layer down, onto CNT assemblies. As for MEA preparations, retinas were held in place with a membrane filter and stainless steel anchor weights. aCSF was perfused into the main chamber (0.5 ml/min) and carboxygenated directly within the chamber. To test the viability of the tissue, some biohybrids were removed after 72 hours and placed onto an active MEA and they always displayed both spiking activity and LFPs on multiple channels.

### Electron microscopy

For TEM, retinas incubated onto loose CNT discs were lifted off from the SiO_2_ substrate using a piece of nitrocellulose membrane filter, halved along the optic disc, then fixed in 2% Glutaraldehyde (Cacodylate buffered), post-fixed in osmium tetroxide, dehydrated in acetone, and embedded in epoxy resin (TAAB, UK). Semi-thin (2 μm) and ultra-thin (70-90 nm) sections were cut perpendicular to the surface of the retina on an Ultracut E ultra-microtome (Reichert-Jung, Austria, now Leica Microsystems), then stained with either toluidine blue (semi-thin sections) or uranyl acetate and lead citrate (ultra-thin sections).

Semi-thin sections were obtained in levels of 3 series comprising 6 sections to be used for staining to locate the CNT islands in the tissue. Thus, each one of these levels corresponded to a 36 μm-long segment of tissue, preventing the inadvertent full sectioning of a large CNT island.

For scanning EM, retinas incubated onto loose CNT discs were lifted off from the SiO_2_ substrate using a piece of nitrocellulose membrane filter, halved along the optic disc, then fixed in 2% Glutaraldehyde (Sorensons buffered), dehydrated in increasing concentrations of ethanol and critical-point dried (CPD 030, Bal-Tec, Lichtenstein, now Leica Microsystems). The samples were coated with a 15 nm gold film using a Polaron E5550 sputter coater (Polaron equipment Ltd, UK, now Quorum Technologies Ltd) before observation in a Cambridge Stereoscan 240.

### Immunohistochemical labelling of Retinal Sections

Mouse eyes were enucleated, fixed for an hour in 4% paraformaldehyde, cryo-protected in a 30% sucrose phosphate buffered saline (PBS) solution for 24 hours, embedded in Tissue-Tek OCT Compound (Sakura, Japan) at −30°C and sectioned in 10 μm slices with a cryostat (HM560, Microm International). Sections were gathered on gelatine-coated slides, air-dried for 24 hours, immersed in de-ionised H_2_O for 1 minute to dissolve the OCT compound and air-dried for 1 hour before immunohistochemical labelling.

Slides were kept at room temperature (if not specified otherwise) in a humidifying chamber and washed 3 times for 5 minutes in 0.1 M PBS (phosphate buffered saline) between each of the following steps in which reagents were diluted in a solution consisting of 0.1% Triton X 100 (Sigma-Aldrich, UK) and 0.1 M PBS: sections were exposed to 0.6% H_2_O_2_ for 20 min (to remove endogenous peroxidases), 20% blocking serum (Normal Horse Serum, Vector Labs, UK) for an hour, 2.5 µl/ml mouse anti-GFAP and 10% blocking serum for 16 hours at 4°C, 2 µl/ml biotinylated anti-mouse IgG (Vector Labs, UK) for 3 hours, 2 µl/ml horseradish peroxidase-streptavidin (Vector Labs, UK) for 3 hours then finally reacted with H_2_O_2_ and DAB.

Nuclear staining was achieved by dipping slides in a solution of Mayer’s Haematoxylin before dehydrating the tissue in increasing concentrations of ethanol (50%, 70%, 90% then twice 100% for 3 minutes each), 3 minutes in 2 consecutive jars of Histoclear (Fisher Scientific, UK) and finally, coverslipped using Histomount (Fisher Scientific, UK).

### Immunohistochemical labelling of Retinal wholemounts

Retinas were fixed in 4% PFA for 1 hour then washed 3 times for 5 minutes in de-ionised water. These were then placed, RGC layer facing up, on gelatine-coated slides and dried overnight. Slides were kept at room temperature (if not specified otherwise) in a humidifying chamber and washed 3 times for 5 minutes in 0.1 M PBS between each of the following steps in which reagents were diluted in a solution consisting of 0.1% Triton X 100 (Sigma-Aldrich, UK) and 0.1 M PBS: sections were exposed to 20% blocking serum (normal goat serum and normal donkey serum, Sigma Aldrich, UK) for 1 hour, 5 µl/ml primary antibodies (mouse anti-GFAP, Sigma Aldrich; rabbit anti-laminin, ABcam) and 10% blocking serum for 20 hours at 4°C, 2 µl/ml secondary antibodies (FITC donkey anti-mouse and Cy3 goat anti-rabbit, Stratech, UK) for 3 hours. After another PBS wash, slides were coverslipped with hard setting Vectashield (Vector labs, UK), then dried overnight.

### Imaging of immunohistochemically stained retinas

Wholemount retinas were visualised with a Nikon A1R point scanning confocal microscope (Nikon Corporation, Japan) with the RGC layer facing the objective. Tiff stacks were processed in ImageJ (NIH, USA). Peroxidase stained retinal sections were visualised on an Olympus BX60 (Olympus, Japan) microscope with a Zeiss Axiocam HRc camera using Zeiss Axio Vision 3.1 software (Carl Zeiss AG, Germany). Fluorescence-labelled retinal sections were acquired on a Leica DMRA microscope (Leica Microsystems) with a Hamamatsu Orca-ER camera (Hamamatsu Photonics, Japan) using the Axio Vision 4.8 (Carl Zeiss AG, Germany) software.

## Acknowledgements

This work was supported by the EPSRC (ES, DTA studentship to CE), Newcastle Healthcare Charity (ES), BBSRC BB/1023526/1 and BB/F011415/1 (ES), the Wellcome Trust 087223/Z/08/Z and 090196/Z/09/Z (ES, JZ and HK), the Israel Ministry of Science and Technology, the Israel Science Foundation, and the European Research Council funding under the European Community’s Seventh Framework Program (FP7/2007–2013)/ERC grant agreement FUNMANIA-306707 (YH). We thank Kathryn White, Vivian Thompson and Tracey Davey from Newcastle University electron microscopy research services for training, assistance and invaluable insights. We also thank Alex Laude for assistance with confocal microscopy, Gavin Clowry and Claudia Racca for advice with immunohistochemistry as well as Tom Smulders for the use of his cryostat-microtome.

## Authors contributionS

CE, JZ and ES designed the experiments. MD-P and YH designed and manufactured the CNT MEAs and other CNT assemblies used for anatomical studies. CE, JZ, HK and ES performed the experiments. CE, JZ, HK and ES analysed the results. CE and ES wrote the manuscript with input from the other authors.

## COMPETING FINANCIAL INTERESTS

The authors declare no competing financial interests.

